# Collective movement of schooling fish reduces locomotor cost in turbulence

**DOI:** 10.1101/2024.01.18.576168

**Authors:** Yangfan Zhang, Hungtang Ko, Michael Calicchia, Rui Ni, George V. Lauder

## Abstract

The ecological and evolutionary benefits of collective behaviours are rooted in the physical principles and physiological mechanisms underpinning animal locomotion. We propose a turbulence sheltering hypothesis that collective movements of fish schools in turbulent flow can reduce the total energetic cost of locomotion by shielding individuals from the perturbation of chaotic turbulent eddies. We test this hypothesis by quantifying energetics and kinematics in schools of giant danio (*Devario aequipinnatus*) compared to solitary individuals swimming under control and turbulent conditions over a wide speed range. We discovered that, when swimming at high speeds and high turbulence levels, fish schools reduced their total energy expenditure (TEE, both aerobic and anaerobic energy) by 63–79% compared to solitary fish. Solitary individuals spend ∼25% more kinematic effort (tail beat amplitude*frequency) to swim in turbulence at higher speeds than in control conditions. However, fish schools swimming in turbulence reduced their three-dimensional group volume by 41–68% (at higher speeds) and did not alter their kinematic effort compared to control conditions. This substantial energy saving highlighted a ∼261% higher TEE when fish swimming alone in turbulence are compared to swimming in a school. Schooling behaviour could mitigate turbulent disturbances by sheltering fish within schools from the eddies of sufficient kinetic energy that can disrupt the locomotor gaits. Providing a more desirable internal hydrodynamic environment could be one of the ecological drivers underlying collective behaviours in a dense fluid environment.

**One-Sentence Summary:** The collective movement of fish schools substantially reduces the energetic cost of locomotion in turbulence compared to that of swimming alone.

## Introduction

Nearly all animal species live with ubiquitous turbulent air or water in nature (1) (2) (3) (4). Hence, turbulent flows affect many aspects of animal biology that are fundamental to lifetime fitness, including dispersal and spawning, the cost of moving for both regional locomotion and long-distance migration, and the dynamics of predator-prey interactions (5). In particular, chaotic turbulent flows (6) (7) (8) directly subject solitary individuals to unpredictable fluid fields and alter body kinematics. For animal species that routinely interact with ambient flow and perceive their fluid environment, when visual input on incoming turbulent flow is limited, individual animals may have limited sensory input and less anticipatory time to adjust body motion on short time scales. Under such challenging conditions, individual animals may have few options for mitigating the energetic costs of living and moving in turbulence.

This challenge is especially formidable for aquatic life in natural channels of rivers and coastal seas (9) (10) (11) (12), because water is 50 times more viscous than air (13) (14) (5) (15) and exerts larger perturbing forces on fish. Moving in turbulence is particularly challenging and energetically expensive for solitary fish. Solitary creek chub (*Semotilus atromaculatus*) swimming in turbulence reduced maximum sustained swimming speed (by 22%) because large turbulent eddies (∼76% of body length) disrupt the movement trajectories of fish (14). Also, the cost of locomotion by solitary Atlantic salmon (*Salmo salar*) can increase by ∼150% in turbulence (16). Studies on animal locomotion and turbulence have profound implications for a better understanding of the planetary ecosystem, e.g., turbulence generated by groups of fish can contribute to vertical mixing of the ocean (17) (18) (19). Despite the widespread interest in understanding how fish interact with turbulence (5) (14) (15) (18) (20) (21) (22) (23) (24) (25) (26) (19) (27) (21) and the ubiquitous interactions of animals and their turbulent fluid environments, no previous study has investigated the effects of turbulent flow on fish schools. This can be due to the complexity of turbulent flow and the dynamic nature of collective animal motion. Could the collective movement of fish schools mitigate the effects of turbulent flow by alternating their locomotor characteristics?

Fish schools could modulate oncoming turbulent flow through the coordination of nearby individuals. Nearly all fish could modify the local fluid environment through vortices shed by their undulatory body motion, and by acting as nearby solid surfaces (28) (29) (30) (31) (32). A school of fishes could reduce the intensity and length scale of oncoming turbulent eddies within the school (Fig. 1). Hence, we propose a “turbulence sheltering hypothesis” that a fish school can shield individuals within the group from ambient turbulence (Fig. 1). If this hypothesis holds, a key prediction is that collective movement reduces the total energy expenditure per unit of biomass compared to that of a solitary individual under the same flow conditions. This study focuses on testing this hypothesis experimentally.

**Fig. 1.**
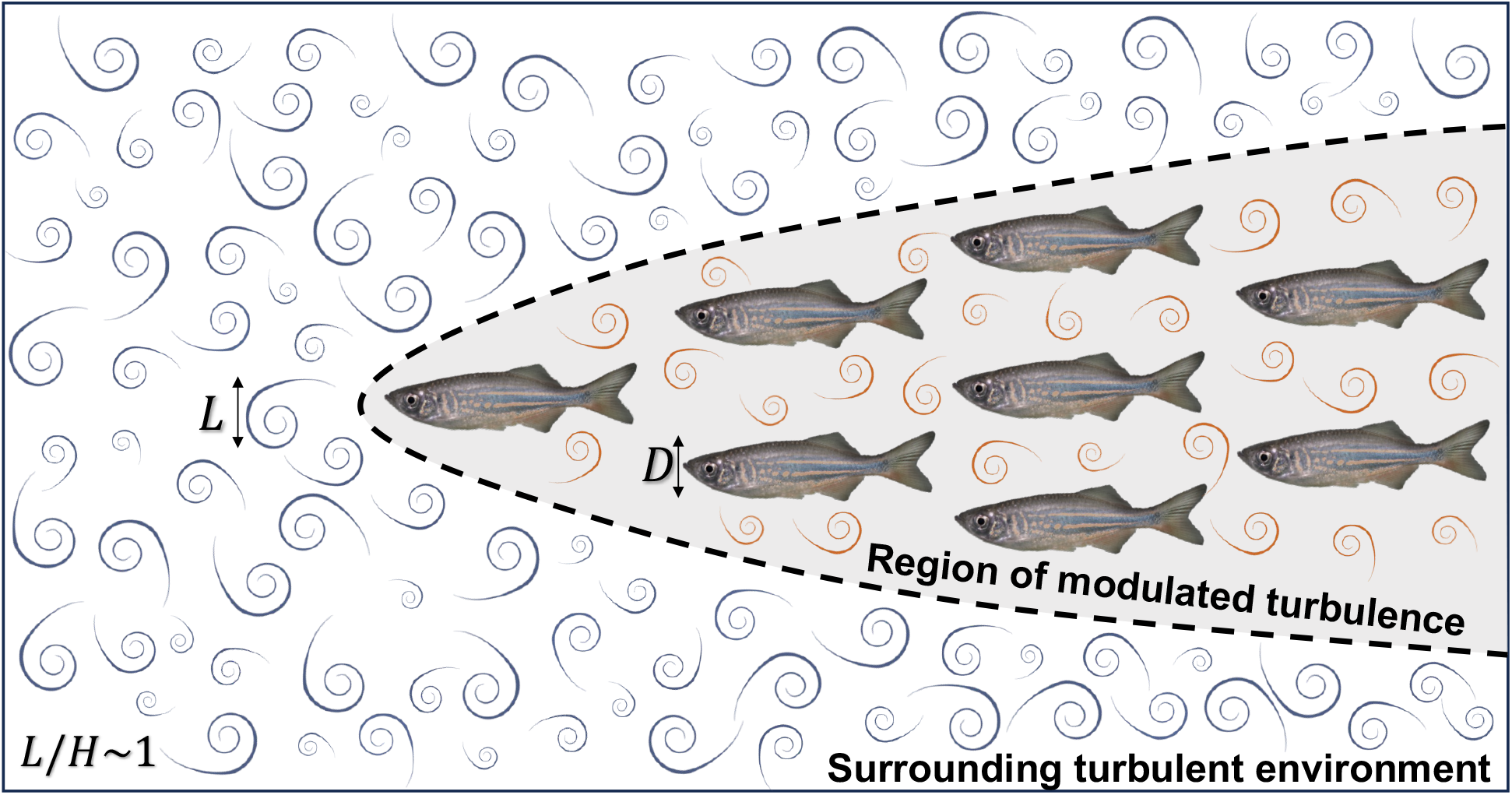
Illustration of the environmental turbulence sheltering hypothesis. Schematic diagram of a school of giant danio (*Devario aequipinnatus*) swimming in oncoming turbulence where the largest eddies have an integral length scale (*L*) on the same order of magnitude as the body depth (D) of the fish. Fish within the school could benefit from a region of reduced turbulence created within the school as a result of nearby neighbours and undulatory body motion modifying flow within the school compared to free stream oncoming flow. As a result, we propose a “turbulence sheltering” hypothesis that fish schools can protect individuals within the group from free-stream turbulence. As a result, we predict fish swimming in turbulence could reduce their locomotor costs by schooling in contrast to swimming alone.

Vertebrates use both aerobic and non-aerobic metabolic energy to support their total energy expenditure (TEE) during locomotion. Aerobic metabolism primarily supports energy use at slower and steady locomotion, while glycolytic metabolism supplies faster and unsteady state high-speed movement (33). Not only do the physiological mechanisms underpinning locomotion shift with speed, but fluid drag also scales as the square of fluid velocity. Hence, increased swimming speeds physically require substantially more metabolic energy. The characterization of a locomotor performance curve (TEE as a function of speed) (34) under both turbulent and control conditions will test the working hypothesis that fish schools could mitigate the expected increase in the energetic cost of moving in turbulence. To test this hypothesis, we directly quantified the locomotor performance curve for both solitary individuals and schools of eight giant danio (*Devario aequipinnatus*) across a wide range of speeds from 0.3 to 8 body lengths sec^-1^. We measured whole-animal aerobic energy expenditure (oxidative phosphorylation) during swimming, as well as excess post-exercise O_2_ consumption (EPOC) to quantify non-aerobic energy expenditure after swimming (high-energy phosphates and substrate-level phosphorylation) (35) (36) (37). In addition, we simultaneously quantified the kinematics of individual fish and those within schools, and measured three-dimensional school volumes to characterize how fish responded to both control and turbulent flow environment.

## Results

### Hydrodynamics of turbulent conditions

Turbulent flows (generated by a passive turbulence grid) exhibit strong fluctuations and chaotic patterns (Fig. 2 A,B) in contrast to controlled flows (generated by a flow straightener). Quantitatively, turbulent testing conditions showed sustainably greater maximum velocity (*F*_1,107_ = 401.9, *p* < 0.001, Fig. 2C), maximum vorticity (*F*_1,107_ = 167.8, *p* < 0.001, Fig. 2D), and maximum (Fig. 2E) and sum shear strength (*F*_1,107_ ≥ 153.7, *p* < 0.001, Fig. 2F). Since turbulence was generated by passing the flow through a passive grid, greater turbulence was reached at higher mean flows; similar rise of values of all these parameters in control flow condition was also observed, but with a greatly reduced rate of increase with velocity (Fig. 2). In addition, the probability density function (p.d.f) of flow velocity along the swimming direction (*u*) and perpendicular direction (*v*) showed the broadening of p.d.f (Fig. 3A & B), a clear indication of intensified turbulence as speed increases. Since turbulence fluctuation velocity increases approximately linearly with the mean flow speed (Fig. 3C), the resulting turbulence intensity, defined as the ratio between the two, remains nearly constant over the flow speeds studied.

**Fig. 2.**
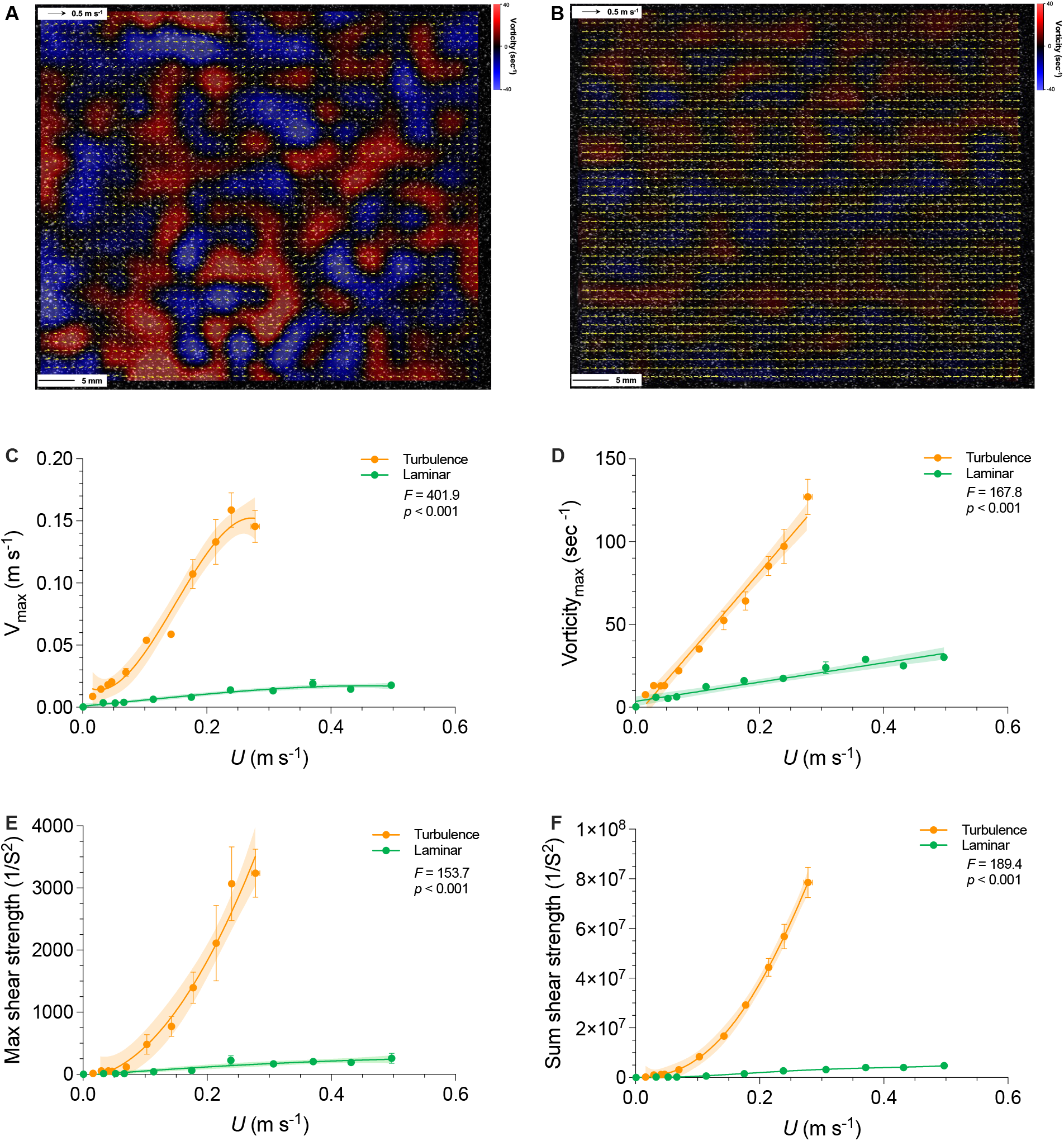
Characterization of hydrodynamic features in controlled and turbulent flows across a range of speeds used for danio schooling energetic and kinematic measurements. Representative flow patterns of (**A**) turbulent and (**B**) control conditions in the swim-tunnel respirometer as quantified by particle image velocimetry. Velocity vectors are yellow arrows, and the vorticity field is shown by blue (-40 sec^-1^) to red (40 sec^-1^) gradient heat maps in the background. The hydrodynamic features of turbulent (orange colour) and controlled (green colour) flows are characterized by (**C**) maximum vertical velocity, (**D**) maximum vorticity, (**E**) maximum and (**F**) the sum of shear strength as a function of absolute speed (meter sec^-1^). More detailed flow characteristics are illustrated in the Supplemental materials for characteristics of the turbulent flow generated in the respirometer (Fig. S2. The statistics in each panel denote the main effect of the flow condition. Shading indicates the 95% confidence interval. Statistical details are available in the statistical analyses section. The controlled fluid condition is a laminarized flow environment.

**Fig. 3.**
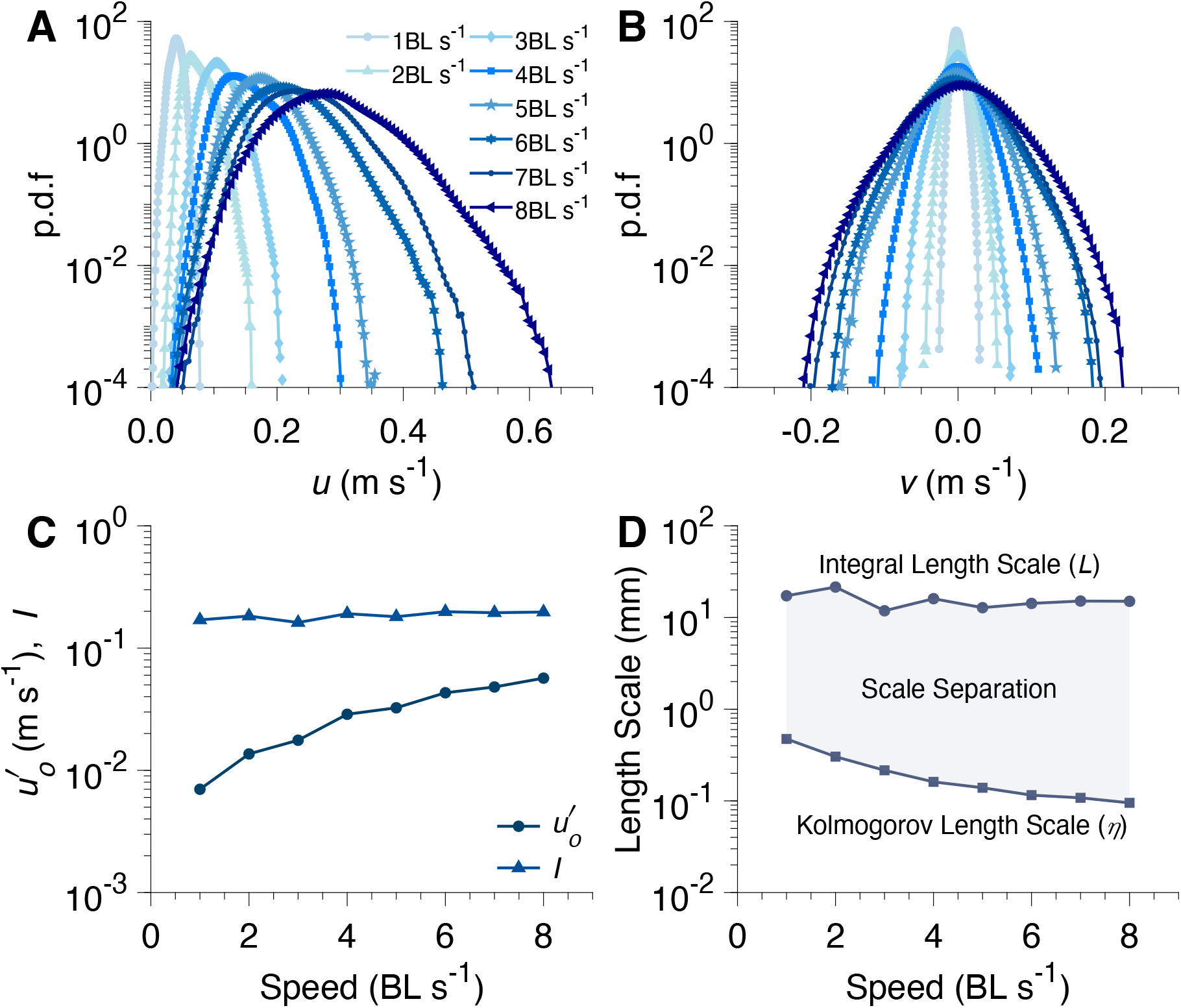
Analysis of turbulent flow in the respirometer used for testing schooling energetics. Probability density function (p.d.f) of velocity components **(A)** parallel (**A**) and **(B)** perpendicular (*v*) to the swimming direction for different **swimming speeds** with the passive turbulence grid. As speed increases, the width of the distribution increases, signifying increasing turbulence. **(C)** The turbulence intensity (*I*) and fluctuation velocity (**A**.) as a function of swimming speed. Fluctuation velocity increases with increasing speed. However, the increase in the fluctuation velocity is proportional to the increase in the speed resulting in a near constant turbulence intensity. (D) Distribution of eddy sizes present for a given swimming speed, from largest (integral length scale) to smallest (Kolmogorov length scale). The eddy size distribution was determined by approximating the energy dissipation rate via computation of the two-dimensional structure functions (See Supplemental Material for further information). The largest eddies were roughly the same size as the fish’s body depth, or 30% of their body length. The control fluid test condition is a laminarized flow environment.

Eddies of various sizes can differentially impact the energetics of fish locomotion. The undulatory motion of fish can respond to eddies smaller than fish size, whereas eddies comparable to the body size may have sufficient energy to change the fish’s movement trajectory, which could result in increased energy expenditure. However, turbulence is notorious for its wide spectrum of scales, which can be quantified by the largest (integral scale) and the smallest (Kolmogorov scale), shown as two separate lines with the range in between showing the full range (Fig. 3D). Based on the energy cascade framework of turbulence eddy scale, the large eddies at the integral length scale (L) were comparable to the fish body depth (D) and are capable of energetically challenging fish locomotion (Fig. 1, 3).

### Energetics of collective movement

We discovered that aerobic metabolic rate – speed curves of fish schools and solitary individuals both are concave upward in turbulence across the entire 0.3–7 BL s^-1^ speed range (Fig. 4). We demonstrate that turbulent flow has resulted in upshifted aerobic metabolic rate – speed curves of solitary individuals across the entire speed range (*F*_1, 69_=4.45, *p*=0.0003) compared to locomotion by fish schools. In particular, the energetic cost of swimming was 32% lower at 6 BL s^-1^ in schools compared to individuals swimming alone in turbulence (719.3 vs 1055 mg O_2_ kg^-1^ h^-1^, *F*_1,69_ *=* 4.1*, p* = 0.0015, Fig. 4A). Aerobic energy conservation enabled by schooling dynamics is more pronounced at the higher speeds when the energy demands are at a premium.

**Fig. 4.**
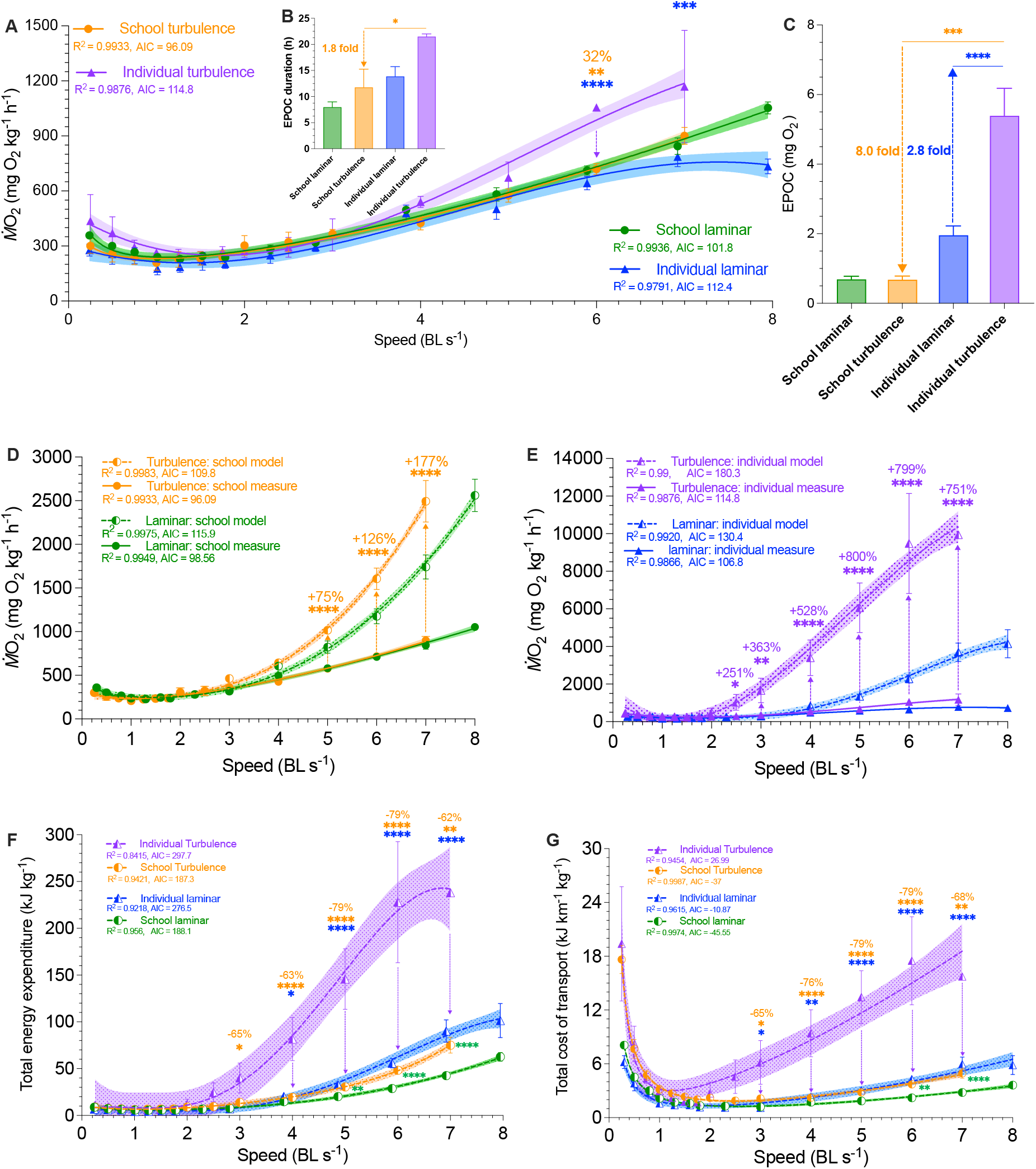
Measurements of aerobic and non-aerobic locomotor costs for fish schools and solitary fish in control and turbulent flow conditions. (**A**) Comparison of concave upward metabolic rate (ṀO_2_)-speed curve over 0.3–8 body length s^-1^ (BL s^-1^) range for fish schools and solitary fish swimming in control and turbulent flow conditions. (**B**) recovery time of excess post-exercise O_2_ consumption (EPOC) and (**C**) EPOC of fish schools and solitary fish after swimming in control and turbulent conditions. Concave upward total ṀO_2_-speed curve of (**D**) fish schools and (**E**) solitary fish when swimming in control and turbulent conditions. The total ṀO_2_-speed curve (dashed line) is calculated from a model that integrates the measurements (solid line) of aerobic and non-aerobic locomotion costs (*see* Supplemental Material). (**F**) Total energy expenditure (TEE) and (**G**) concave upward total cost of the transport (TCOT)-speed curves for fish schools and solitary fish when swimming in control and turbulent conditions. Total Energy Expenditure (TEE) and the Total Cost of transport (TCOT) are calculated using the sum of aerobic and non-aerobic costs. Green colour = fish schools in control conditions (n=5); blue colour = solitary fish in control conditions (n=5); orange colour = fish schools in turbulence (n=4); purple colour = solitary fish in turbulence (n=3). Statistical significance is denoted by asterisk(s). Shading indicates the 95% confidence interval. Statistical details are available in the statistical analyses section. The control fluid condition is a laminarized flow environment.

Because fish body musculature operating at high frequencies during locomotion at high speed mostly uses white muscle fibres powered in part through glycolysis (38) (39), we predict that the schooling dynamics should also conserve non-aerobic energy (estimated by EPOC) and reduce the recovery time when compared with solitary individuals. Indeed, the non-aerobic cost for fish schools to swim through the entire speed range in turbulence was nearly 8-fold lower than that of solitary individuals (EPOC: 0.68 vs 5.4 mg O_2_*, t=*7.0*, p=*0.0005, Fig. 4C). Fish that swam in schools recovered 1.8-fold faster than solitary individuals (EPOC duration: 11.8 vs 21.5 h, *t=*2.3*, p=*0.035).

As a result, both total energetic expenditure (TEE) and total cost of transport (TCOT) of fish schools was 62–79% lower than that of solitary individuals swimming in turbulence over the 3–7 BL s^-1^ range Fig. 4G). Non-aerobic costs contribute 72–83 % of total energy consumption in solitary fish, whereas the non-aerobic contribution was only 20–40% in fish schools (Table S1). If non-aerobic locomotor costs are not accounted for, 75–177% of locomotor energy expenditure in solitary individuals (Fig. 4D), and 251–800% of the energy expenditure in fish schools (Fig. 4E) would not have been accounted for.

Despite the substantial energy saving enabled by fish schooling in turbulence compared to swimming in the same turbulent environment alone, fish schools do not completely eliminate the effects of turbulence on locomotor cost. After accounting for *both* aerobic and non-aerobic energy costs, we discovered that the TEE of fish schools swimming in turbulence was 51–76% higher (*F*_1, 7_*=*54.3, *p* ≤ 0.0072, Fig. 4F) than the cost for fish schools swimming under control (low turbulence) hydrodynamic conditions over the speed range of 5–7 BL s^-1^, and TCOT of fish schools was 68 and 76% higher in turbulence than in control low-turbulence flow (*F*_1, 91_=36.2, *p* ≤ 0.007) at 6 and 7 BL s^-1^ respectively (Fig. 4G). The proportional increase in the total energetic cost of swimming in turbulence is again mostly from non-aerobic energy production. Nevertheless, the magnitude of the increased total costs of swimming in turbulence is only a fraction (9–24%) of the costs for solitary individuals. Solitary individuals spent 190–342% higher total energy swimming in turbulence across the speed range of 2.5–7 BL s^-1^ compared to locomotion in control fluid conditions (Fig. 4F, G).

### Kinematics

Using high-speed videography, we discovered that, as speed increases to speeds >1.7 BL s^-1^, school volume becomes smaller, and the 3-D convex hull representing school volume is reduced by 74% as individuals within the school swim in closer proximity (*F*_12,143_ *=* 11.74, *p* < 0.0001). Over the speed range of 1.7–7 BL s^-1^, fish schools become denser and form a prolate spheroid shape compared to the streamlined school structure when fish schools swim in controlled flows of the same speed. The 3-D convex hull volume of fish schools swimming in turbulent conditions is 40–68% lower than that for schools swimming in control conditions at similar speeds (*t*_1,8_ *≥* 2.147, *p* ≤ 0.032, Fig. 5).

**Fig. 5.**
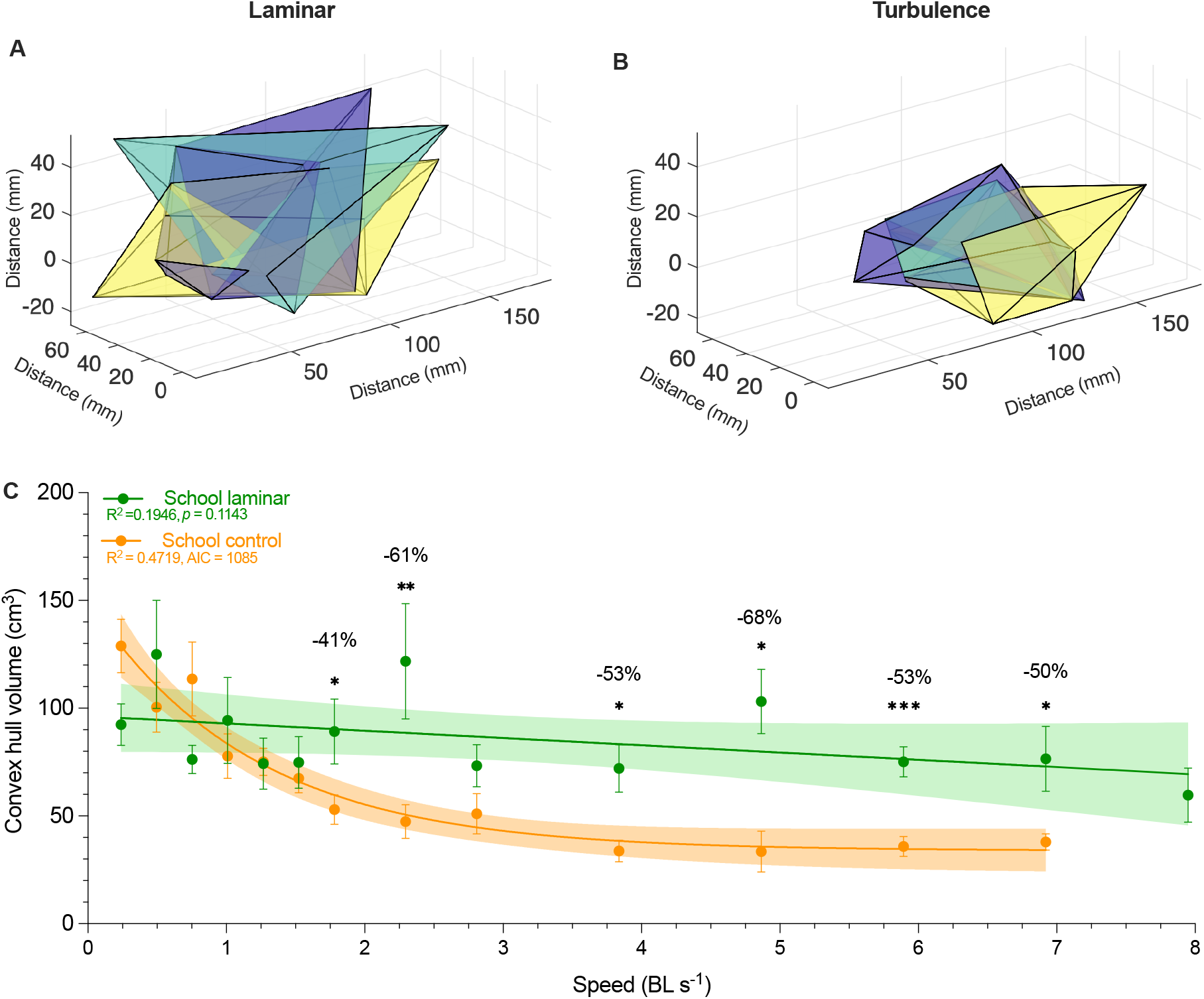
Characterization of fish school three-dimensional volume in turbulent and control flow conditions. Representative three-dimensional (3-D) convex hull volume of fish schools in (**A**) control and (**B**) turbulent flow conditions. (**C**) 3-D convex hull volume as a function of speed for fish schools swimming in control (n=9 snapshots per speed increment) and turbulent (n=12 snapshots per speed increment) flow conditions. Statistical significance is denoted by asterisk(s). Shading indicates the 95% confidence interval. Statistical details are available in the statistical analyses section. The control fluid condition is a laminarized flow environment.

Also, kinematics of individual fish (Tail beat frequency, *f*_tail_; tail beat amplitude, Amp_tail_; *f*_tail_•Amp_tail_) within schools swimming in turbulence were not different from when fish schools swam in control flow conditions (Fig. 6F, MANOVA: *F*_1,25_ ≤ 0.028, *p* ≥ 0.868). However, solitary fish spend up to 22% more effort (estimated as tail beat frequency times amplitude, *f*•Amp_tail_) swimming in turbulence than for control flow locomotion (Fig. 6E, MANOVA: *F*_1,28_ ≥ 1.167, *p* ≤ 0.006). Solitary fish increase *f*_tail_ at lower speeds (ANOVA: *F*_1,575_ ≥ 4.67, *p* ≤ 0.031, Fig. 6A) and also reducing Amp_tail_ (ANOVA: *F*_1,639_ ≥ 26.35, *p* < 0.001, Fig. 6C). As a result, swimming effort remains the same at lower speeds when compared to solitary individuals swimming in controlled flows, as indicated by both kinematics (*f*•Amp) (MANOVA: *F*_1,28_ = 1.167, *p* = 0.281) and energetics (Fig. 6F, ANOVA: *F*_1,71_ ≤ 1.23, *p* ≥ 0.96). As speed increases in turbulence, solitary fish increase Amp_tail_ by 26% (Fig. 6C) (ANOVA: *F*_14,639_ = 3.69, *p* < 0.001), and there is no further increase in *f*_tail_ (Fig. 6A). Locomotor effort (*F*_tail_•Amp_tail_) increased with speed at a greater rate for individuals in turbulence compared to individuals swimming under control conditions over the range of speeds tested (ANOVA: *F*_7,536_ = 173.14 *p* ≤ 0.006, Fig. 6E).

**Fig. 6.**
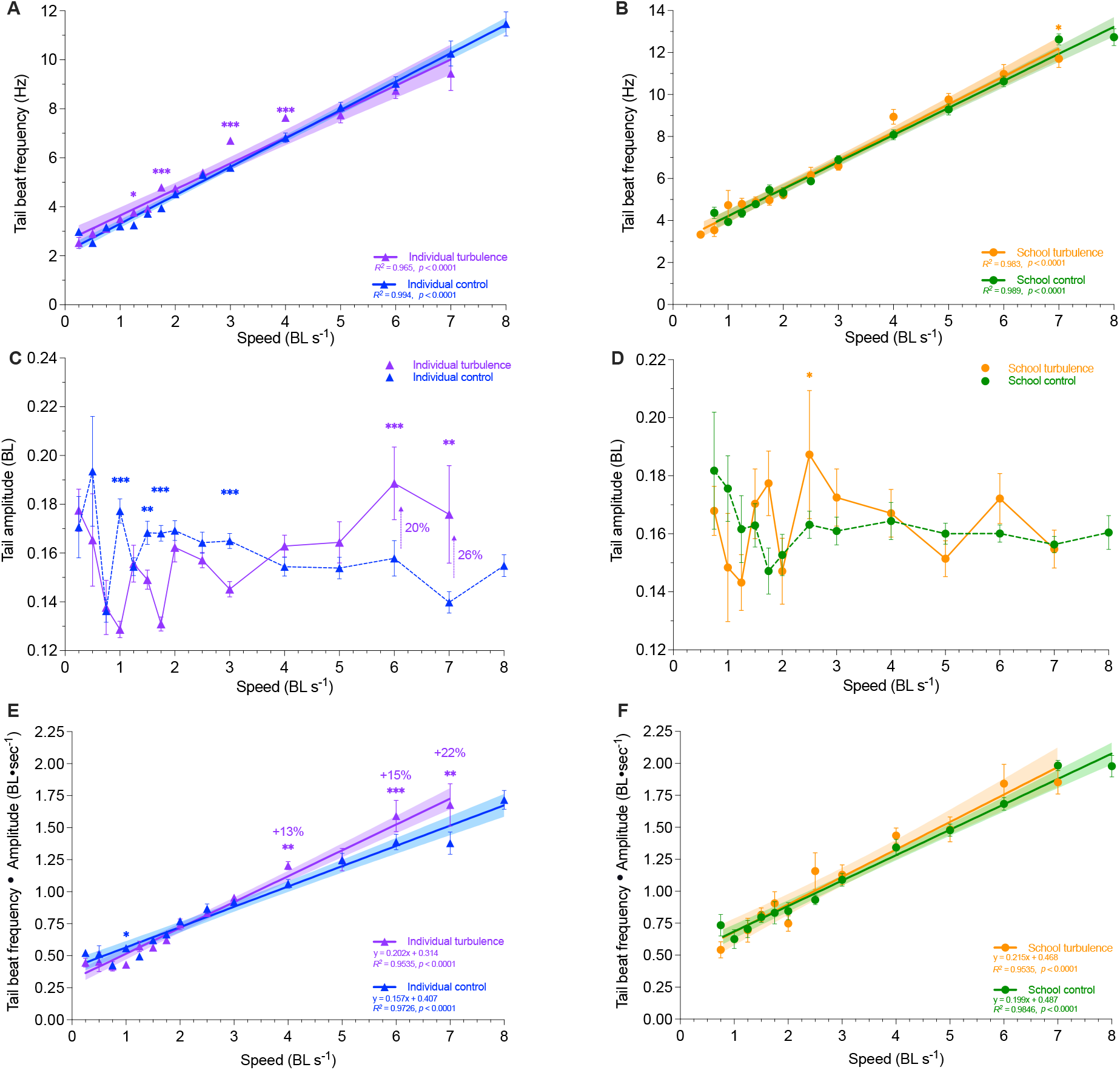
Kinematic data on individual fish within the school during locomotion in control and turbulent flow conditions. Tail beat frequency (*f*_tail_) of (**A**) solitary fish and (**B**) fish schools across 0.3–8 body length s^-1^ (BL s^-1^) in control and turbulent flow conditions. Tail beat amplitude (Amp_tail_) of (**C**) solitary fish and (**D**) fish schools across 0.3–8 body length s^-1^ (BL s^-1^) in control and turbulent flow conditions. An estimate of swimming effort, *F_t_*_ail_ • Amp_tail_, of (**E**) solitary fish and (**F**) fish schools across 0.3–8 body length s^-1^ (BL s^-1^) in control and turbulent flow conditions. Green colour = fish schools in control conditions (n=295–379 sequences); blue colour = solitary fish in control conditions (n=351–416 sequences); orange colour = fish schools in turbulence (n=104–146 sequences); purple colour = solitary fish in turbulence (n=220–258 sequences). Statistical significance is denoted by asterisk(s). Shading indicates the 95% confidence interval. Statistical details are available in the statistical analyses section. The control fluid condition is a laminarized flow environment.

## Discussion

Despite the ubiquity of vertebrates moving in environments with turbulent flows, we know remarkably little about the energetic costs of moving in turbulence as a collective group compared to moving in the same conditions as a solitary individual. Hence, we integrate three lines of evidence (energetics, individual kinematics, and schooling dynamics) and compare them between fish schools and solitary individuals over the same speed range in turbulence.

Our results support the turbulence sheltering hypothesis. We first discovered that the collective movement of fish schools substantially dampens the effects of turbulence by downshifting the locomotor performance curve at higher speeds, compared to when solitary individuals swim in turbulence. Fish schools in turbulence expend up to 79% less energy than fish under control conditions (Fig. 4F). One of the essential mechanisms by which fish schools dampen turbulent disturbances is by collectively swimming in up to a 68% tighter schooling formation compared to control conditions. As a result, fish swimming within schools showed no difference in their swimming kinematics regardless of whether the fish schools swim in turbulence or control conditions. We also discovered that one of the key reasons for higher locomotor costs when solitary individuals swim in turbulence is the increase in tail beat amplitude (Amp_tail_) at higher speeds (Fig. 6C), which increases the kinematic effort of swimming as estimated by tail beat frequency times amplitude (*f*•Amp_tail_) (Fig. 6E, F). As a result, the total energy expenditure (TEE) and total cost of transport for solitary individuals, including both the aerobic and non-aerobic energy contributions, are ∼261% higher compared to when solitary individuals swim in control conditions (Fig. 4F, G).

Therefore, collective behaviour provides effective turbulence sheltering, not only mitigating the kinematic responses needed in turbulent flow, but also providing a large energetic advantage by downshifting most of the locomotor performance curve. We highlight three key considerations regarding collective movement in turbulent flows: 1) How does collective movement dampen the turbulent disturbance on locomotor energetics? 2) How do body kinematic patterns act as a linchpin between fluid dynamic and energetic effects? 3) How does the hydrodynamic scale of turbulence relative to fish size matter for broader considerations of fish locomotor ecology?

### 1) Schooling dynamics dampens the effects of turbulence

Fish schools (*i.e.* giant danio) are effective at reducing the energetic costs of swimming in turbulence. The TEE and TCOT of solitary individuals are 188–378% higher than that of fish schools at higher speeds, per kilogram of biomass. By studying both aerobic and non-aerobic locomotor costs, we discovered that most of the energy saving stems from the reduced use of non-aerobic energy (Table S1 & S2). Specifically, fish schools swimming in turbulence generated an 8-fold *lower* excess post-exercise oxygen consumption (EPOC, including the use of high-energy phosphate stores and glycolytic energy contributions) than that of a solitary individual. The aerobic cost of swimming at higher speeds was also 47% higher in solitary individuals compared to that of fish schools. Collectively, over the entire range of swimming up to maximum and sustained speeds, schooling dynamics effectively dampens the additional metabolic costs of moving in the turbulent flow by 3.8 folds (the integral area under the TEE curve of schools versus individuals in turbulence: 165.5 vs. 633.3 kj kg^-1^).

Solitary individuals increased tail beat frequency (*f*_tail_) but reduced tail beat amplitude (Amp_tail_) to compensate for turbulent disturbances at the lower speeds (≤ 43% critical swimming speed, *U*_crit_), which yielded the same kinematic effort (estimated as *f*_tail•_Amp_tail_) as swimming under control flow conditions. As water velocity increases to ≥ 86% of *U*_crit_, solitary individuals can no longer increase *f*_tail_ to compensate for the effects of turbulence. Instead, solitary individuals increased Amp_tail_ to compensate for the turbulent disturbances which increased kinematic effort and reflected in substantially higher energetic cost. In contrast, schooling is highly effective at reducing the effects of turbulent eddies on fish kinematics within the school, and fish within a school in turbulence move similarly to fish swimming in controlled flow.

Bioenergetics is critical to understanding the cost of behaviours (40) (41) (42), and allows direct quantification of the amount of energy used to answer fundamental questions including “How much does a behaviour cost?” and “How do altering environmental conditions affect this cost?”. Energetic measurements are particularly useful in evaluating hypotheses involving locomotion occurring over a range of movement speeds and in comparison to control conditions. The kinematic effort of fish swimming (estimated as *f*•Amp_tail_) suggested that energy saving by schooling danio in turbulence is ∼25%, whereas direct measurement of energy expenditure shows a total energy saving of ∼79%. Kinematic analyses typically use snapshots of body motion of individuals to quantify biomechanical effort, but do not necessarily reflect total energy use, particularly when sampled intermittently during an incremental speed test. Our energetic measurements detected that turbulent flow increased the total cost of locomotion of fish schools by 51–76% at higher speeds compared to when fish schools swim in controlled flow conditions, whereas fish schools swimming in turbulent and control conditions showed no difference in tail kinematics. Although understanding the biomechanics of locomotion and movement are essential adjuncts to studying locomotion energetics, the cost of movement for any animal is also governed by the underlying physiological mechanisms relating to the cardiorespiratory system, circulatory system, musculature and metabolic pathways that generate the ATP needed for movement. We thus advocate here for integrated studies of kinematics and energetics, and caution that kinematic studies alone may not reflect actual levels of energy use.

### 2) Kinematics and schooling dynamics in turbulence

To better understand the interactions between fish kinematics and fluid dynamics, we characterized Strouhal number (St, dimensionless undulatory propulsive effort at a movement speed) (43) (44) and Reynolds number (Re, dimensionless fluid speed showing the ratio between inertial and viscous forces) (13). Regardless of whether solitary individuals or fish schools swim in controlled or turbulent conditions, the St ranged 0.25–0.35 (Fig. 7A). The general relationship of St and Re is independent of added turbulence and whether or not fish swim within a school.

**Fig. 7.**
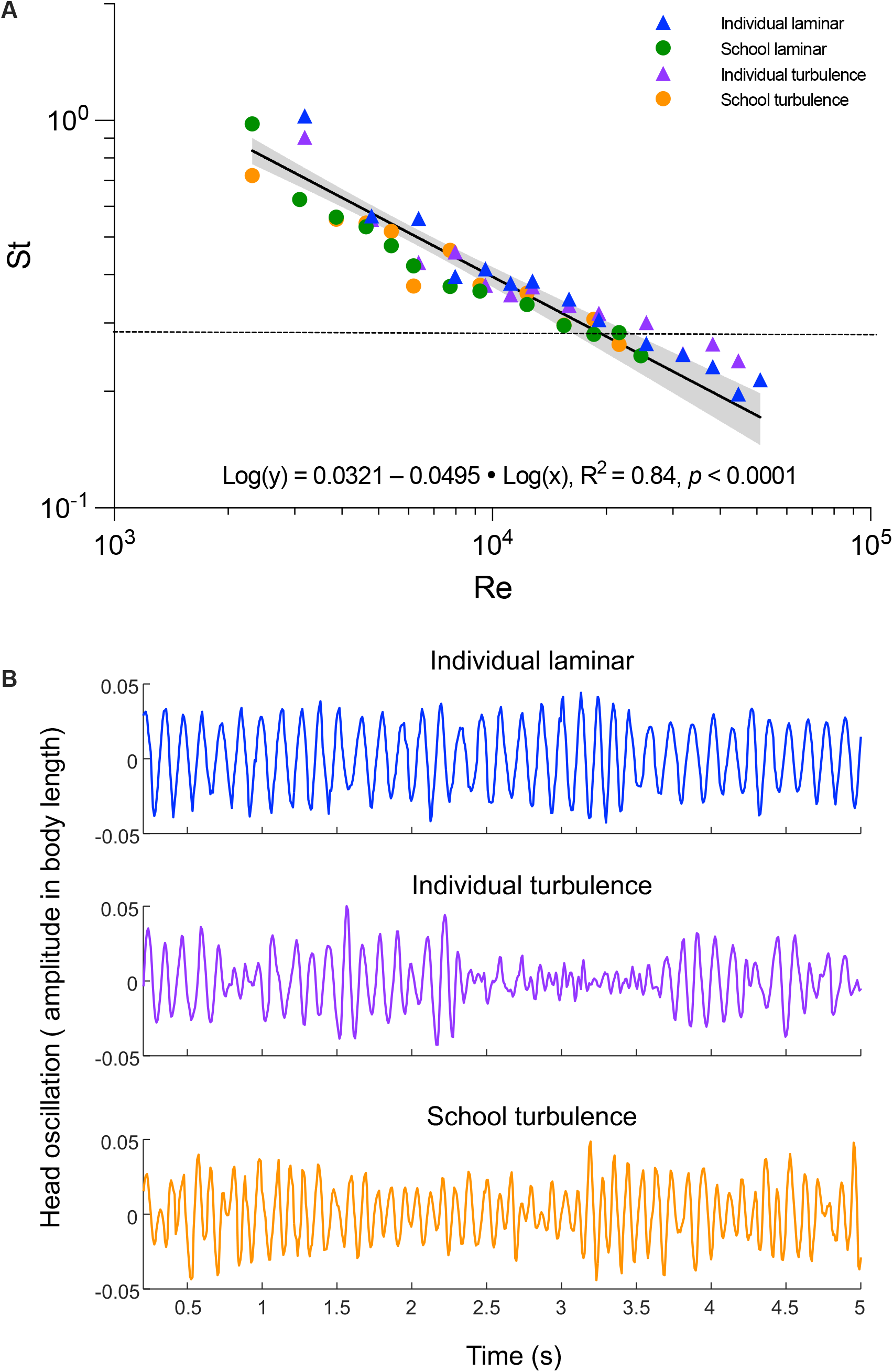
Responses of schooling fish to the fluid dynamic environment within fish schools under control and turbulent conditions. (**A**) Relationship between two key dimensionless parameters (Strouhal number and Reynolds number) for fish swimming in schools and alone under both control and turbulent conditions. Locomotion follows a generally linear relationship (R^2^=0.84, *p*<0.0001). The horizontal dashed line provides the hypothesized relationship of St and Re in the turbulent flow following (45). The St number at the lowest speed, when fish exhibited unsteady turning behaviour, is excluded from the analysis to prevent bias in the scaling relationship of directional oscillatory propulsion. (**B**) Plots of fish head lateral oscillation through time (blue: individual laminar, purple: individual turbulence, orange: school turbulence) in water velocity of 6 body length sec^-1^. Shading indicates the 95% confidence interval. Statistical details are available in the statistical analyses section. The control fluid condition is a laminarized flow environment.

The general relationship of Re and St (the log-log linear relationship, Fig. 7A) falls in the vicinity of a general scaling hypothesis for aquatic undulatory locomotion (45). However, these data show that even in the turbulent flows giant danios maintain the linear relationship between St and Re instead of transitioning to a different relationship as proposed by the previous scaling hypothesis (45). It is possible that, if the Re of locomotion is increased beyond Re > 10^5^ (*e.g*. larger species moving at faster speeds), the relationship of Re and St could change, but our data do not support the previously suggested scaling of locomotor St and Re in turbulence. Future laboratory studies are needed to better inform the relationship of Re and St across a wide range of swimming velocities in turbulent conditions that are ubiquitous in nature.

Since our data collapse onto a single scaling relationship (Fig. 7A), how can we explain the difference in metabolic rates between turbulent and control conditions? As a completed swim oscillatory body wave typically starts from the head of the fish, hence we tracked the oscillation at the nostril region of the fish head (oscillatory amplitude as a function of time) to examine the fluid conditions surrounding the fish. We reason that locations within a school can reduce the turbulent disturbances that would otherwise increase locomotor costs for solitary fish. Individuals swimming alone under controlled conditions show a regular pattern of head oscillation frequency and near-constant oscillation amplitude (Fig. 7B, blue wave). However, the head oscillations of solitary individuals in turbulent flows are irregular and of varying frequency (Fig. 7B, purple wave). Individuals within a school swimming in turbulence have rhythmical head oscillations and more distinct peaks than those under turbulent conditions (Fig. 7B, orange wave), and the pattern of head oscillation is more like that of solitary individuals swimming in control flow conditions. The benefit of swimming within a school is most likely the result of the tighter schooling formation as fish schooling volumes are reduced in turbulent flows. Smaller inter-individual distances allow the myriad of hydrodynamic mechanisms that are associated with reduced cost of locomotion to become effective, including fish swimming side-by-side, in front and behind other fish, and in the reduced velocity zone behind two fish (34) (46) (32) (47) (48). These results suggest that fish schools act as effective “shelters” that enhance hydrodynamic mechanisms that reduce locomotor cost, a strategy that is not available to individual fish swimming alone in turbulence.

As a topic for future investigation, we suggest that the fish schools could alter the size scale of turbulent eddies within the school and thus reduce the impact that environmental perturbations have on the cost of individuals swimming. Fish schools could function as band-pass filters as a result of their undulatory body motion and the proximity of individuals within the school to each other. This collective undulatory movement could create a more predictable flow field within the group that reduces kinematic efforts and swimming costs compared to individuals swimming alone in turbulence.

Measuring flow conditions within a school would provide a more direct and detailed understanding of how turbulent eddies within fish schools are modified in comparison to the free-stream turbulent flow field. However, this is currently a considerable experimental and technical challenge. Even when using multiple laser light sheets or volumetric approaches to illuminate flow within a school, the bodies of fish in a dense school inevitably cast shadows and hinder the resolution of imaging turbulent flow within the school. In the absence of direct measurements for within-school flow fields, kinematic and energetic data provide the best available evidence for the turbulence sheltering hypothesis.

### 3) Turbulence length scale and fish ecology

Turbulent disturbances on animal movement are multifaceted and context-specific. Turbulent flows are chaotic and unpredictable by fish, and contain energy over a wide spectral range (Figs. 2, 3), which differs from a Kármán vortex wake where a solitary fish can interact with a regular and predictable pattern of oncoming vortices (49). Fish swimming in a Kármán vortex street can save energy by tuning their body dynamics to interact with oncoming vortices, and greatly reduce their muscle activity and energetic cost (3) (50). When fish are exposed to the true chaotic turbulent flow with a length scale on the same order of magnitude of body size, as in the experiments presented here, the energetic cost of locomotion greatly increases.

Demonstrating the reduction in the total cost of locomotion in turbulence by group dynamics in aquatic vertebrates has direct implications for the movement ecology of migratory species. For example, a “feast-or-famine” life history is common to many migratory species. Food availability can be scarce and fluctuate during the migration journey (51). Migratory (fish) species typically accumulate energy and nutrients during the feeding season prior to undertaking a long migration. As a result, migratory species often rely on a finite number of onboard stores of metabolic substrates to fuel migration. We demonstrated here that the total cost of transport when fish schools move through turbulence is substantially decreased. Thus, fish schools should migrate a *longer* distance for the same amount of energy. By migrating in a collective group, as many fish species do, fish should be able to sustain migration against unexpected costs associated with changing environmental stressors such as heat, hypoxic episodes, and storms.

Fish encounter turbulence not only in natural environments as a result of rapid stream flows, bottom topography, or obstacles in the water, but also under conditions where human-designed structures such as dams or fish passage structures create turbulence (52) (53) (54). A key issue in considering how fish must contend with these structures is understanding the length scale of turbulence encountered by fishes and whether or not fish prefer and could even benefit from a turbulent environment (55). Our experiments utilized a passive turbulence grid to generate turbulent eddies with a length scale approximating the body depth of the giant danio studied. In nature, however, turbulent flows can differ in the turbulence length scale, and in the energy present at each eddy size, and variations in turbulence scale can be important for understanding the effects of turbulence on animals of different sizes. It remains unknown if fish (either individuals or schools) select particular turbulent length scales when swimming that could also allow locomotor energy savings in contrast to the increased costs demonstrated at other length scales. Perhaps the design of fish passage devices for habitat restoration should consider the ratio of turbulence eddy scale relative to the animal size to improve locomotor ability and reduce the cost of movement for fish (52). Alternatively, energetically costly turbulence flow generators could serve as aquatic barriers to perturb invasive species.

Given the ubiquity of turbulent flows in natural aquatic ecosystems, we suggest that one of the important roles of collective behaviour in fish species is to shelter individuals within a collective group from challenging hydrodynamic conditions. More broadly, our study proposes that vertebrate collectives can also function as a larger size biological entity than the solitary individual, which could reduce the effect of turbulence perturbation on animal movement. Using aquatic vertebrates moving in the dense water fluid as a model system to directly demonstrate energy saving can be the foundation for future studies of the ‘turbulence sheltering’ hypothesis in flying and terrestrial vertebrates. Locomotor performance curves, where metabolic or kinematic variables are evaluated against swimming speed, are a useful comparative framework to broadly understand the energetic cost of collective movement (34).

## Materials and Methods

### Experimental animals

Experiments were performed on giant danio (*Devario aequipinnatus*) that were acquired from a local commercial supplier near Boston, Massachusetts USA. Five schooling groups are randomly distributed and housed separately in five 37.9 l aquaria (n=8 per tank). The five solitary individuals are housed separately in five 9.5 l aquaria (n=1 per tank). All aquaria have self-contained thermal control (28 °C), an aeration system (>95 % air saturation, % sat.) and a filtration system. Water changes (up to 50% exchange ratio) were carried out weekly. Fish were fed *ad libitum* daily (TetraMin, Germany). Animal holding and experimental procedures were approved by the Harvard Animal Care IACUC Committee (protocol number 20-03-3).

### Experimental system – Integrated Biomechanics & Bioenergetic Assessment System (IBAS)

The experimental system and similar experimental protocols are available in (56). To promote reproducibility, we reiterate the methodologies in detail and the additional experimental detail specific to the study of collective movement in turbulent conditions.

The core of our experimental system is a 9.35-l (respirometry volume plus tubing) customized Loligo® swim-tunnel respirometer (Tjele, Denmark). The respirometer has an electric motor, and a sealed shaft attached to a propeller located inside the respirometer. By regulating the revolutions per minute (RPM) of the motor, the water velocity of the motor can be controlled.

The swim-tunnel respirometer is oval-shaped. The central hollow space of the respirometry increases the turning radius of the water current. As a result, the water velocity passing the cross-section of the swimming section (80 × 80 × 225 mm) is more homogenous (validated by PIV). Moreover, a honeycomb flow straightener (80 × 80 × 145 mm) is installed in upstream of the swimming section to create laminar flow (validated by PIV). The linear regression equation between RPM and water velocity (V) of control flow is established (V = 0.06169•RPM – 5.128, R^2^ = 0.9988, *p <* 0.0001) by velocity field measured by particle image velocimetry (PIV).

To increase the signal-to-noise ratio for the measurement of water dissolved O_2_, a water homogenous loop is installed 95 cm downstream of the propeller and the water is returned to the respirometer 240 cm before the swimming section. The flow in the water homogenous loop moves (designated in-line circulation pump, Universal 600, EHEIM GmbH & Co KG, Deizisau, Germany) in the same direction as the water flow in the swimming tunnel. A high-resolution fibre optic O_2_ probe (Robust oxygen probe OXROB2, PyroScience GmbH, Aachen, Germany) is sealed in the homogenous loop at downstream of the circulation pump (better mixing) to continuously measure the dissolved O_2_ level in the water (recording frequency ∼1 Hz, response time < 15s). The oxygen probe was calibrated to anoxic (0 % sat., a solution created by super-saturated sodium sulphite and bubbling nitrogen gas) and fully aerated water (100 % sat.). The background ṀO_2_ in the swim-tunnel respirometer was measured for 20 min before and after each trial. The average background ṀO_2_ (< 6% of fish ṀO_2_) was used to correct for the ṀO_2_ of fish. The pre-filtered water (laboratory grade filtration system) is constantly disinfected by UV light (JUP-01, SunSun, China) located in an external water reservoir to suppress the growth of microbial. Water changes of 60% total volume occurred every other day and a complete disinfection by sodium hypochlorite is conducted weekly (Performance bleach, Clorox & 1000 ppm).

To simultaneously measure schooling dynamics and swimming kinematics, the customized oval-shaped swim-tunnel respirometer is located on a platform with an open window beneath the swimming section. The platform is elevated 243 mm above the base to allow a front surface mirror to be installed at a 45° angle. This mirror allows a high-speed camera (FASTCAM Mini AX50 type 170K-M-16GB, Phontron Inc., United States, lens: Nikon 50mm F1.2, Japan) to record the ventral view. The second camera (FASTCAM Mini AX50 type 170K-M-16GB, Phontron Inc., United States, lens: Nikon 50mm F1.2, Japan) is positioned 515 mm to the side of the swimming section to record a lateral view. Synchronized lateral and ventral video recordings were made at 125 fps, and each frame was 1024 by 1024 pixels. To avoid light refraction passing through the water and distorting the video recordings, the swim-tunnel respirometry is not submerged in the water bath. Temperature regulation of the respirometer is achieved by regulating room temperature, installing thermal insulation layers on the respirometry and replenishing the water inside the respirometer from a thermally regulated (28 °C, heater: ETH 300, Hydor, United States & chiller: AL-160, Baoshishan, China) water reservoir (insulated 37.9-l aquarium) located externally.

The aerated (100% sat., air pump: whisper AP 300, Tetra, China) reservoir water is flushed (pump: Universal 2400, EHEIM GmbH & Co KG, Deizisau, Germany) to the respirometer through an in-line computer-controlled motorized ball valve (U.S. Solid) installed at the in-flow tube. The other in-line one-way valve is installed at the out-flow tube. The out-flow tube is also equipped with a valve. The value is shut during the measurement period, a precautionary practice to eliminate the exchange of water between the respirometer and the external reservoir when the water moves at a high velocity inside the respirometer. This flushing was manually controlled to maintain DO above 80 % sat. Every time the respirometer was closed to measure ṀO_2_, the water temperature fluctuates no more than 0.2 °C. The water temperature inside the respirometer is measured by a needle temperature probe (Shielded dipping probe, PyroScience GmbH, Aachen, Germany) sealed through a tight rubber port of the respirometer.

To allow fish to reach the undisturbed quiescent state during the trial, the entire Integrated Biomechanics & Bioenergetic Assessment Platform (IBAP) is covered by laser blackout sheet (Nylon Fabric with Polyurethane Coating; Thorlabs Inc, New Jersey, United States). The room lights are shut off and foot traffic around the experimental rig is restrained to the absolute minimum. Fish are orientated by dual small anterior spots of white light (lowest light intensity, Model 1177, Cambridge Instruments Inc, New York, United States) for orientation (one to the top and the other to the side) of the swimming section. The test section is illuminated by infrared light arrays.

### Creating turbulent flows in swim-tunnel respirometer

We used a passive turbulence grid (height × width: 7.5 × 8.6 cm) to generate the turbulence flow for the swimming section in the swim-tunnel respirometer (Fig. S1). The turbulence grid has a configuration of 3 × 3 square openings (each opening is in 1.5 × 1.5 cm). The openings produce 9 streams of jets which mix and form turbulent flow. The turbulent grid is upstream of the swimming section (Fig. S1). The opening of the grid is guarded by thin metal wires to prevent fish from going through. The turbulence grid is effective in generating turbulence, as illustrated by the fluid dynamic features of the turbulences measured by PIV (*see* Fig. S2 and Figs. 2 and 3). As a result, the linear regression equation between RPM and average water velocity (V) of turbulent flow is changed (V = 0.03515•RPM – 1.597, R^2^ = 0.9985, *p <* 0.0001) and quantified by velocity field measured by particle image velocimetry (PIV) (*see* Fig. S3). Quantifying average water velocity allowed us to match the swimming kinematics and metabolic energy consumption of tested fish at the same mean speed between control and turbulent flows.

### Experimental Protocol

The same individuals or schools are repeatedly measured in laminar or turbulent flow to control for biological variations. Giant danio (*Devario aequipinnatus*) is a model species, capable of actively and directionally swimming from a minimum to maximum sustained speeds (0.3–8.0 body lengths s^-1^; BL s^-1^ & Reynolds number range of 6.4•10^3^ to 1.8•10^5^ in controlled flow). We studied five replicate schools and five replicate individuals drawn from within each school. Swimming performance test trials were conducted with *Devario aequipinnatus* fasted for 24 hours, a sufficient period for a small-sized species at 28 °C (*i.e.* high resting ṀO_2_) to reach an absorptive state. In fact, we observed no specific dynamic action, in the amount of oxygen consumed for digestion during the first diurnal cycle (Fig. S3). Prior to the swimming performance test, testing fish were gently weighted and placed in the swim-tunnel respirometer. The fish swam at 35% *U*_crit_ for 30 mins to help oxidize the inevitable but minor lactate accumulation during the prior handling and help fish become accustomed to the flow conditions in the swim-tunnel respirometer (57). After this time, the fish to be tested were habituated (>20 hours) to the respirometer environment under quiescent and undisturbed conditions. During this time, we used an automatic system to measure the resting ṀO_2_ for at least 19 hours. Relays (Cleware GmbH, Schleswig, Germany) and software (AquaResp v.3, Denmark) were used to control the intermittent flushing of the respirometer with fresh water throughout the trial to ensure O_2_ saturation of the respirometer water. ṀO_2_ was calculated from the continuously recorded dissolved O_2_ level (at 1 Hz) inside the respirometer chamber. The intermittent flow of water into the respirometer occurred over 930 s cycles with 30 s where water was flushed into the respirometer and 900 s where the pumps were off and the respirometer was a closed system. The first 240 s after each time the flushing pump was turned off were not used to measure ṀO_2_ to allow O_2_ levels inside the respirometer to stabilize. The remaining 660 s when the pumps were off during the cycle were used to measure ṀO_2_. The in-line circulation pump for water in the O_2_ measurement loop stayed on throughout the trial.

We characterize the locomotor performance curve of fish using an established incremental step-wise critical swimming speed (*U*_crit_) test (35). The first preliminary trial determined the *U*_crit_ of this population of *Devario aequipinnatus* as 8 BL s^-1^. Characterizing the swimming performance curve required a second preliminary trial to strategically select 10 water velocities (0.3, 0.5, 0.8, 1.0, 1.3, 1.5, 1.8, 2.3, 2.8 BL s^-1^) to bracket the hypothesized concave upward metabolism-speed curve at the lower speed (< 40% *U*_crit_). Additional five water velocities (3.8, 4.9, 5.9, 6.9, 8.0 BL s^-1^) are used to characterize the exponentially increasing curve to the maximum and sustained swimming speed, *U*_crit_ (*see* Fig. S5). Altogether, 14 points provide a reliable resolution to characterize the locomotor performance curve. At each water velocity, fish swam for 10 mins (58) to reach a steady state in ṀO_2_ at low speeds (*see* Fig. S6). Above 40% *U*_crit_, ṀO_2_ can become more variable (59). Hence, in this protocol, we focus on measuring the sustained aerobic energy expenditure by calculating the average ṀO_2_ for each 10-min velocity step using Eqn 1. The respirometry system reaches a stable signal-to-noise ratio once the sampling window is longer than 1.67 mins (*see* Fig. S7), well within the duration of the velocity step to obtain a stable signal-to-noise ratio for calculating ṀO_2_ (59). At the 5^th^ min of each velocity step, both ventral and lateral-view cameras are triggered simultaneously to record 10-sec footage at 125 frames per second, at 1/1000 shutter speed and 1024 ×1024 pixel resolution. Thus, both data streams of ṀO_2_ and high-speed videos are recorded simultaneously. The *U*_crit_ test is terminated when 12.5% of fish in the school or a solitary individual touches the back grid of the swimming section for more than 20 secs (57). The *U*_crit_ test lasted ∼140 mins and estimates the aerobic portion of energy expenditure over the entire range of swimming performance.

To measure the contribution of non-aerobic O_2_ cost, where most of the cost is related to substrate-level phosphorylation, and to calculate the total energy expenditure for swimming over the entire speed range, we measured excess post-exercise oxygen consumption (EPOC) after the *U*_crit_ test for the ensuing 19 hours, recorded by an automatic system. Most previous measurements of EPOC after *U*_crit_ test have used a duration of ∼5 hours, but our extended measurement period ensured that longer duration recovery O_2_ consumption (EPOC) was measured completely as fish were exercised to *U*_crit_ (*see* summary table in *16*). The intermittent flow of water into the respirometer occurred over 30 s to replenish the dissolved O_2_ level to ∼95% sat. For the following 900 s the flushing pump remained closed, and the respirometer became a closed system, with the first 240 s to allow O_2_ saturation inside the respirometer to stabilize. The remaining 660 s when the flushing pump was off during the cycle were used to measure ṀO_2_ (*see* Eqn 1). The cycle is automated by computer software (AquaResp v.3) and provided 74 measurements of ṀO_2_ to compute EPOC. Upon the completion of the three-day protocol, the school or individual fish are returned to the home aquarium for recovery. The fish condition was closely monitored during the first 48 hours after the experiment, during which no mortality was observed.

### Bioenergetic measurement and modeling

To estimate the steady-rate whole-animal aerobic metabolic rate, ṀO_2_ values were calculated from the sequential interval regression algorithm (Eqn. 1) using the dissolved O_2_ (DO) points continuously sampled (∼1 Hz) from the respirometer.

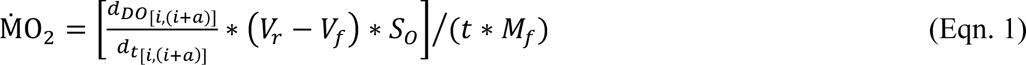

Where ^*d_Do_*^/*d_t_* is the change in O_2_ saturation with time, *V_r_* is the respirometer volume, *V_f_* is the fish volume (1 g body mass = 1 ml water), *S_o_* is the water solubility of O_2_ (calculated by AquaResp v.3 software) at the experimental temperature, salinity and atmospheric pressure, *t* is a time constant of 3600 s h^-1^, *M_f_* is fish mass, and *a* is the sampling window duration, *i* is the next PO_2_ sample after the preceding sampling window.

To account for allometric scaling, the ṀO_2_ values of solitary fish were transformed to match the size of the individual fish in the school using an allometric scaling exponent (b = 0.7546). The calculation of the scaling relationship [Log_10_(ṀO_2_) = b•Log10(*M*) + Log10(*a*), where *M* is the body mass & *a* is a constant] was performed by least squares linear regression analysis (y = 0.7546•x + 0.2046; R^2^ = 0.6727, *p* < 0.0001) on the 180 data points of metabolic rate and body mass from a closely related species (the best available dataset to our knowledge) (61). The allometrically scaled ṀO_2_ values were used to derive other energetic metrics (listed below) for the solitary fish. The energetic metrics of fish schools are calculated from the mass-specific ṀO_2_.

The resting oxygen uptake (ṀO_2rest_), the minimum resting metabolic demands of a group of fish or a solitary individual, is calculated from a quantile 20% algorithm (62) using the ṀO_2_ estimated between the 10^th^–18^th^ hour and beyond the 32^nd^ hour of the trial. These are the periods of quiescent state when fish completed the EPOC from handling and swimming test.

The excess post-exercise oxygen consumption (EPOC) is an integral area of ṀO_2_ measured during post-exercise recovery, from the end of *U*_crit_ until reached ṀO_2rest_ plus 10% (60). This approach reduces the likelihood of overestimating EPOC due to spontaneous activities (60). To account for the allometric scaling effect, we used the total amount of O_2_ consumed (mg O_2_) by the standardized body mass of fish (1.66 g) for fish schools and solitary fish.

We model EPOC (*i.e.* non-aerobic O_2_ cost) to estimate a *total* O_2_ cost over the duration of the swimming performance test. Our conceptual approach was pioneered by Brett (35) in fish and is also used in sports science (33). Mathematical modeling was applied to study the effects of temperature on the cost of swimming for migratory salmon (57). We improved the mathematical modeling by applying the following physiological and physics criteria. The first criterion is that significant accumulation of glycolytic end-product occurred when fish swimming above 50% *U*_crit_ (63) which corresponds to > ∼40% ṀO_2max_ (or ∼ 50% aerobic scope) (33). This is also when fish start unsteady-state burst-&-glide swimming gait (63). The second criterion is that the integral area for the non-aerobic O_2_ cost during swimming can only differ by ≤ 0.2% when compared to EPOC. The non-aerobic O_2_ cost during swimming is the area bounded by modeled ṀO_2_ and measured ṀO_2_ as a function of time when fish swim > 50% *U*_crit_ (*see* Fig. 2A & Table S2). The third criterion is that total energy expenditure is expected to increase exponentially with swimming speed (Fig. S7). Specifically, these curves were fitted by power series or polynomial models, the same models that describe the relationship between water velocity and total power and energy cost of transport (Fig. S7). Following these criteria, the non-aerobic O_2_ cost at each swimming speed is computed by a percentage (%) modifier based on the aerobic O_2_ cost (Table S1 & S2). The exponential curve of total O_2_ cost as swimming speed of each fish school or solitary individual was derived by an iterative process until the difference between non-aerobic O_2_ cost and EPOC met the 2^nd^ criterion. The sum of non-aerobic O_2_ cost and aerobic cost gives the total O_2_ cost.

The best model fitting following the relationships between water velocity and energetic costs of locomotion suggests that the glycolysis starts at 2–3 BL s^-1^ for fish swimming in the turbulence flow, whereas the same model suggests that the glycolysis starts at 4–5 BL s^-1^ for fish swimming in the controlled flow. We are confident in this model, because the engaging glycolysis at the lower swimming speed for the fish swimming in turbulent flow corresponded with their lower *U*_crit_ (controlled flow: 8 BL s^-1^ vs. turbulent flow: 7 BL s^-1^). The model of aerobic and non-aerobic energy costs of locomotion enabled the estimation of total energy expenditure and total cost of transport as detailed below:

Total energy expenditure (TEE) is calculated by converting total O_2_ cost to kJ × kg^-1^ using an oxy-calorific equivalent of 3.25 cal per 1 mg O_2_ (64).

Total cost of transport (COT), in kJ × km^-1^ × kg^-1^ is calculated by dividing TEE by speed (in km × h^-1^) (58).

### Hydrodynamic flow visualization & analysis

We used a horizontal plane of laser sheet (LD pumped all-solid-state 532nm green laser, 5w, MGL-N-532A, Opto Engine LLC) to visualize and quantify the hydrodynamic conditions of laminar and turbulent flows (*i.e.* particle image velocimetry, PIV). The horizontal planes of the water fluid field in laminar and turbulent conditions were calculated from consecutive video frames (1024 × 1024 pixels) using DaVis v8.3.1 (LaVision Inc., Göttingen, Germany). A vector field (covering a horizontal plane of 34.3 cm^2^ with 2193 vectors) is characterized by a sequential cross-correlation algorithm applied with an initial interrogation window size of 64 × 64 pixels that ended at 12 × 12 pixels (3 passes, overlap 50%). To optimize the signal-to-noise ratio on the sequential cross-correlation algorithm, we used an increment of 10 frames at the lowest speed of laminar and turbulent flow conditions. The rest of the speeds in both flow conditions were analyzed using an increment of 1 frame. Five sets of consecutive video frames (1^st^, 250^th^, 500^th^, 750^th^ and 990^th^ frame out of 1000 frames) in laminar (n=5) and turbulent flows (n=5) were used for calculating the metrics.

To optimize the sampling resolution of consecutive PIV frames, the six lowest testing speeds (106, 131, 156, 181, 256, 356 RPM) of both flow conditions were captured by 1000 frame rate per sec (shutter speed: 1/1000), whereas the six highest testing speed (456, 556, 656, 756, 856 RPM) of both flow conditions were captured by 2000 frame rate per sec (shutter speed: 1/1000) using a high-speed camera (FASTCAM Mini AX50 type 170K-M-16GB, Phontron Inc., United States, lens: Nikon 50mm F1.2, Japan).

From the fluid field, we used DaVis (v8.3.1) to calculate the following fluid parameters:

V_max_ (m s^-1^): Extract the maximum value of the velocity at the vector that is perpendicular to the free-stream flow.

Maximum vorticity (sec^-1^): calculated according to the central difference scheme with four closest neighbours at the horizontal plane. This method achieves a high spatial resolution.

Maximum shear strength (1/S^2^): maximum value of the shear strength. Shear strength is the positive Eigenvalue of the Matrix for two-dimensional vorticity on the horizontal plane.

Sum of shear strength (1/S^2^): sum of the shear strength. The calculation of shear strength is stated above.

To compute the fluctuation velocity, turbulence intensity, and eddy size distribution in the turbulent flow, a subsection of the full velocity field was extracted. This subsection spans an area of 22.4 cm^2^ and was located approximately 4 cm from the passive grid (Fig. S1) and 0.5 cm from the walls of the tunnel. For this analysis, velocity fields were computed on all 2000 frames. For further information about the quantification of turbulence and the distribution of eddy sizes, readers can refer to the supplementary material (*see* computing fluctuation velocity and turbulence intensity).

### Three-dimensional kinematic data extraction from high-speed videography

We used two synchronized 10-sec high-speed videos (lateral and ventral views, at each speed) for kinematic analyses. We calibrated the field of view of the high-speed cameras using a direct linear transformation for three-dimensional kinematic reconstruction (DLTdv8) (65) by applying a stereo calibration to the swimming section of the respirometer (*see* Fig. S8). We digitized the anatomical landmarks of fish (*see* Fig. S10) to obtain the X, Y, Z coordinates for each marker at the 1^st^ sec, 5^th^ sec and 10^th^ sec for videos recorded at each speed. These coordinates are used to calculate the following kinematic parameters. All the calculations are validated on the known length and angle of test objects inserted into the tank working section.

Strouhal number (St) represents the dimensionless flapping frequency and amplitude at a given movement speed. 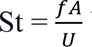 Where *f, A* and *U* are the tailbeat frequency, amplitude, and) swimming speed (13). The measurement is conducted on the calibrated high-speed video in video analysis software (Phontron FASTCAM Viewer 4, Photron USA, Inc.).

Reynolds number (Re) represents the dimensionless fluid inertial to viscous forces at a given speed. 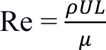, where ρ and μ are the density and dynamic viscosity of the water, and *U* and *L* are the swimming speed and length of the fish (13). Water density and dynamic viscosity are given at 28 °C.

In addition to the manual digitization, we also developed a contrast-based segmentation algorithm for automatic tracking of 2D kinematics in the ventral view. The video analysis was performed in MATLAB (R2022b). We processed each frame in the video independently. We first converted the frame into grayscale and then used a brightness threshold to obtain masks of the fish (Fig S9A). We removed masks with an area larger than that expected for a single fish, excluding masks with multiple overlapping fish. For every streamwise location on the mask (x-location), we calculated the midpoint across the span to obtain the midline of the animal at each instant. The midlines were pieced together across frames according to their locations. We centered and rotated a series of midlines to account for the rigid-body component of the fish body. At this stage of the image analysis, we obtained midline envelopes of the fish (Fig. S9B). We extracted a time series of head and nose spanwise oscillation, identified peaks and troughs of the signal and calculated the amplitude and frequency of the fish (Fig. S9C, D). We excluded time series that are shorter than a tail beat cycle and those associated with unsteady swimming, either with a high relative velocity with flow or fast rotation.

To further characterize the head oscillation of fish in different flow environments, we processed trajectories that are several dozen cycles long at 6 BL s^-1^, extracted through manual digitization (DLTdv8a) (8.2.10) (65). To distinguish the nose oscillation from the background movement of the fish during the period, we performed Fast Fourier Transform (FFT) in MATLAB (R2022b). The FFT analyses enabled us to separate the original time series of the nose trajectories into background motion (low-frequency component) and oscillation due to fish swimming (high-frequency component). The latter is plotted in Fig 7B.

### Statistical analyses

Measurement points are presented as mean ± s.e.m. For the metrics that failed normality tests, logarithm transformations were applied to meet the assumptions of normality of residuals, homoscedasticity of the residuals, and no trend in the explanatory variables. We conducted supervised statistical tests to specifically evaluate our hypotheses about the effects of turbulence on the biomechanics and bioenergetics of fish swimming, either in school or alone. The statistical comparisons for the different responses of fish schools (or solitary fish) between swimming in laminar flows and in turbulent flow used a mixed effects model (laminar flow vs. turbulent flow & swimming speed) with Holm–Šídák *post-hoc* tests. The statistical comparisons for the difference between fish schools and solitary fish swimming in the turbulence used a general linear model (solitary fish vs. fish schools & swimming speed) with Holm–Šídák *post-hoc* tests. The statistical comparison for the characteristics of fluid dynamics between laminar and turbulent flow conditions used a general linear model that used speed as a covariance. The statistical comparison of EPOC between fish schools and solitary fish after performing the *U*_crit_ test in turbulent conditions used an unpaired t-test. The statistical analyses were conducted in SPSS v.28 (SPSS Inc. Chicago, IL, USA). The best-fitting regression analyses were conducted using Prism v.9.4.1 (GraphPad Software, San Diego, CA, USA). 95% C.I. values were presented for all regression models as shaded areas around the regression or data points. Statistical significance is denoted by *, **, ***, **** for *p*-values of ≤ 0.05, ≤ 0.01, ≤ 0.001, ≤ 0.0001 respectively.

## Acknowledgments

Many thanks to members of Lauder Laboratory for numerous discussions about fish schooling behaviour, for comments on the manuscript, to Dr. Robin Thandickal for assistance with the convex hull calculations, and to Cory Hahn for fish care.

## Funding

Funding provided by the National Science Foundation grant 1830881 (GVL), the Office of Naval Research grants N00014-21-1-2661 (GVL), N00014-16-1-2515 (GVL), 00014-22-1-2616 (GVL), and a Postdoctoral Fellowship of Natural Sciences and Engineering Research Council of Canada PDF - 557785 – 2021 (YZ).

## Author contributions

Conceptualization: YZ, GL. RN

Methodology: YZ, GL, HK, MC, HK, RN

Investigation: YZ, HK, MC

Visualization: YZ, GL, MC, HK, RN

Funding acquisition: GL, YZ, RN

Project administration: GL, RN

Supervision: GL, RN

Writing – original draft: YZ

Writing – review & editing: YZ, GL, HK, MC, RN

## Competing interests

Authors declare that they have no competing interests.

## Data and materials availability

All data are available in the main text or the supplementary materials

## Supplementary Materials

Figs. S1 to S9

Supplementary Text

Tables S1 to S2

## Notes

### Competing Interest Statement

The authors have declared no competing interest.

